# Fibroblastic Reticular Cell Response to Dendritic Cells Requires Coordinated Activity of Podoplanin, CD44 and CD9

**DOI:** 10.1101/793141

**Authors:** Charlotte M. de Winde, Spyridon Makris, Lindsey Millward, Jesús Cantoral Rebordinos, Agnesska C. Benjamin, Víctor G. Martínez, Sophie E. Acton

## Abstract

Lymph node expansion is pivotal for adaptive immunity. CLEC-2^+^ migratory dendritic cells (DCs) interact with fibroblastic reticular cells (FRCs) to inhibit podoplanin-dependent actomyosin contractility, permitting FRC spreading and lymph node expansion. However, the molecular mechanisms controlling lymph node remodelling are not fully understood. We asked how podoplanin is regulated on FRCs in the early phase of lymph node expansion *in vivo*, and further, which other FRC markers are required for FRCs to respond to CLEC-2^+^ DCs. We find that expression of podoplanin and its partner proteins CD44 and CD9 in FRCs is coregulated by CLEC-2, and is differentially expressed by specific lymph node stromal populations *in vivo*. We find that beyond contractility, podoplanin is required for polarity and alignment of FRCs. Both CD44 and CD9 act to dampen podoplanin-dependent contractility, and colocalize with podoplanin in different areas of the cell membrane. Independently of podoplanin, CD44 and CD9 affect the degree of cell-cell contact and overlap between neighbouring FRCs. Further, we show that both CD44 and CD9 are required for FRCs to spread and form protrusions in response to DCs. Our data show that remodelling of the FRC cytoskeleton is a two-step process requiring podoplanin partner proteins CD44 and CD9. Firstly, CLEC-2/podoplanin-binding drives relaxation of actomyosin contractility, and secondly FRCs form protrusions and spread which requires both CD44 and CD9. Together, we show a multi-faceted response of FRCs to DCs, which requires CD44 and CD9 in addition to podoplanin.

## Introduction

During the adaptive immune response, the lymph node rapidly expands to accommodate the increased number of proliferating lymphocytes (1–3). Effective immune responses require lymph node remodelling at speed, while maintaining tissue architecture, but it is equally important that remodelling is reversible, and that tissue damage is prevented.

The lymph node is a highly structured organ consisting of different functional zones, which are organised by a connected network of fibroblastic reticular cells (FRCs) (4). T-cell zone FRCs (TRCs) must endure and adapt to the pressure of expanding T-cell populations rapidly requiring additional space (1–3). Throughout expansion, the FRC network remains connected and aligned with extracellular matrix structures (2, 3, 5). Initially, FRCs elongate to stretch the existing network, before proliferating to expand the lymph node further (2–4). Migratory dendritic cells (DCs), expressing the C-type lectin-like receptor CLEC-2, are required to initiate FRC network remodelling (2, 3). When CLEC-2 expression is deleted in DCs, lymph nodes fail to expand relative to controls and the FRC network becomes disrupted (2, 3). CLEC-2 interacts with the glycoprotein podoplanin on the FRC network (2, 3, 6). Podoplanin connects the cytoskeleton to the cell membrane through ERM (Ezrin-Radixin-Moesin) binding and drives activation of RhoA/C (2, 3, 7). CLEC-2^+^ migratory DCs bind and cluster podoplanin, uncoupling podoplanin from RhoA/C activity, and permitting rapid lymph node expansion (2, 3). However, the molecular role of podoplanin on FRCs, and its requirement for FRCs to respond to CLEC-2^+^ migratory DCs, is incompletely understood.

Podoplanin was discovered almost simultaneously in a wide variety of tissues and cell types, and has therefore been assigned multiple names (podoplanin, gp38, Aggrus, PA2.26, D2-40, T1*α*) based on its function in different contexts (8). Podoplanin is widely expressed and plays a pivotal role in the correct development of heart, lungs, secondary lymphoid tissues and lymphatic vasculature. Podoplanin *null* mice exhibit embryonic lethality due to cardiovascular problems or die shortly after birth of respiratory failure (9–11), and exhibit defective blood-lymphatic vasculature separation (12, 13). Podoplanin expression by FRCs is essential for lymph node development, and the maintenance of high-endothelial venule function via interactions with CLEC-2-expressing platelets (14, 15). Podoplanin has a very short cytoplasmic tail (16), and the two serine residues can be phosphorylated, which regulates cell motility (17). The cytoplasmic tail of podoplanin consists of only nine amino acids (16), therefore it is suggested that podoplanin would require partner proteins to execute its functions. Many partner proteins have already been identified (8, 18), but their functions in lymph node remodelling have not been addressed.

Here, we found that two known podoplanin partner proteins, the hyaluronan receptor CD44 (19, 20) and tetraspanin CD9 (21), are transcriptionally regulated in response to CLEC-2, and their expression is controlled on the surface of FRC populations *in vivo* during an immune response. In other biological contexts, CD44 and CD9 link extracellular cues to intracellular signalling. Tetraspanins are a superfamily of four-transmembrane proteins that form tetraspanin-enriched microdomains via interactions with each other and partner proteins. These microdomains spatially organize the plasma membrane into a tetraspanin web, which facilitates cellular communication (22, 23). The interaction of podoplanin with tetraspanin CD9 is mediated by CD9 transmembrane domains 1 and 2, and this interaction impairs cancer metastasis by inhibiting platelet aggregation (21). This is suggestive of a functional role in CLEC-2/podoplanin signalling since platelets express high levels of CLEC-2. CD9 also controls cell fusion and adhesion (24–26). To date, no function of CD9 has been assigned to FRC biology. The interaction of podoplanin with CD44 is mediated by their extracellular domains (19), but it was recently reported to also involve transmembrane and cytosolic regions (20). The podoplanin/CD44 interaction at tumour cell protrusions promotes cancer cell migration (19). Similar to podoplanin, CD44 also binds ERM proteins (27). Interestingly, in NIH/3T3 fibroblasts, coexpression of CD44 and podoplanin reversed the hypercontractile phenotype seen in cells overexpressing podoplanin (2), suggesting an inhibitory function for CD44 in driving actomyosin contractility in fibroblasts. It has previously been shown that podoplanin and CD44 both reside in cholesterol-rich membrane regions on MDCK cells (28). CLEC-2 binding to FRCs drives podoplanin clustering into cholesterol-rich domains (2), but the function of these podoplanin clusters is unknown.

In this study, we seek to further our understanding on the role of podoplanin and its partner proteins in FRC function during lymph node expansion. We investigate the functions of CD44 and CD9 in the response of FRCs to CLEC-2^+^ DCs, the critical initiating step for FRC network spreading and lymph node expansion. We also investigate the requirement for podoplanin signalling with these partner proteins to control FRC functions beyond actomyosin contractility.

## Results

### Increased expression of podoplanin and CD44 on TRCs during adaptive immune responses

It is known that FRC numbers remain constant in the first acute phase of lymph node expansion, and that FRC proliferation is induced later during tissue remodelling (2, 3). In the acute phase, CLEC-2 expressed on DCs inhibits podoplanin-driven actomyosin contractility in FRCs (2, 3), allowing the FRC network to elongate to accommodate the increased number of lymphocytes (1–3). However, the molecular mechanisms are still incompletely understood.

In agreement with previous reports (1, 2), we find that lymph node mass increased 2-3 fold (Fig. 1a) following IFA/OVA immunisation, driven primarily by the increased numbers of lymphocytes (Supplementary Fig. 1). Despite a rapid increase in lymph node mass (Fig. 1a) and total cellularity (Fig. 1b), the number of CD45^-^ lymph node stromal cells do not significantly increase in the first 5 days post-immunization (Fig. 1c). However, stromal cell numbers remain higher than steady state levels, even after lymphocytes have started to traffic out of the tissue (day 7-14) (Fig. 1c and Supplementary Fig. 1). We compared the response of lymphatic endothelial cells (LEC; CD31^+^PDPN^+^), marginal reticular cells (MRC; CD31^-^PDPN^+^MAdCAM-1^+^) and T-cell zone FRCs (TRC; CD31^-^PDPN^+^MAdCAM-1^-^) through the acute phase of lymph node expansion (day 0-7) (Fig. 1d), and found that surface expression of podoplanin does not change on LECs (Fig. 1e), but is increased in both MRCs and TRCs (Fig. 1f-g). This was unexpected, as we hypothesized that podoplanin expression may decrease, since podoplanin-driven actomyosin contractility is inhibited during this early phase of lymph node expansion, due to binding of CLEC-2 on incoming migratory DCs.

**Fig. 1.**
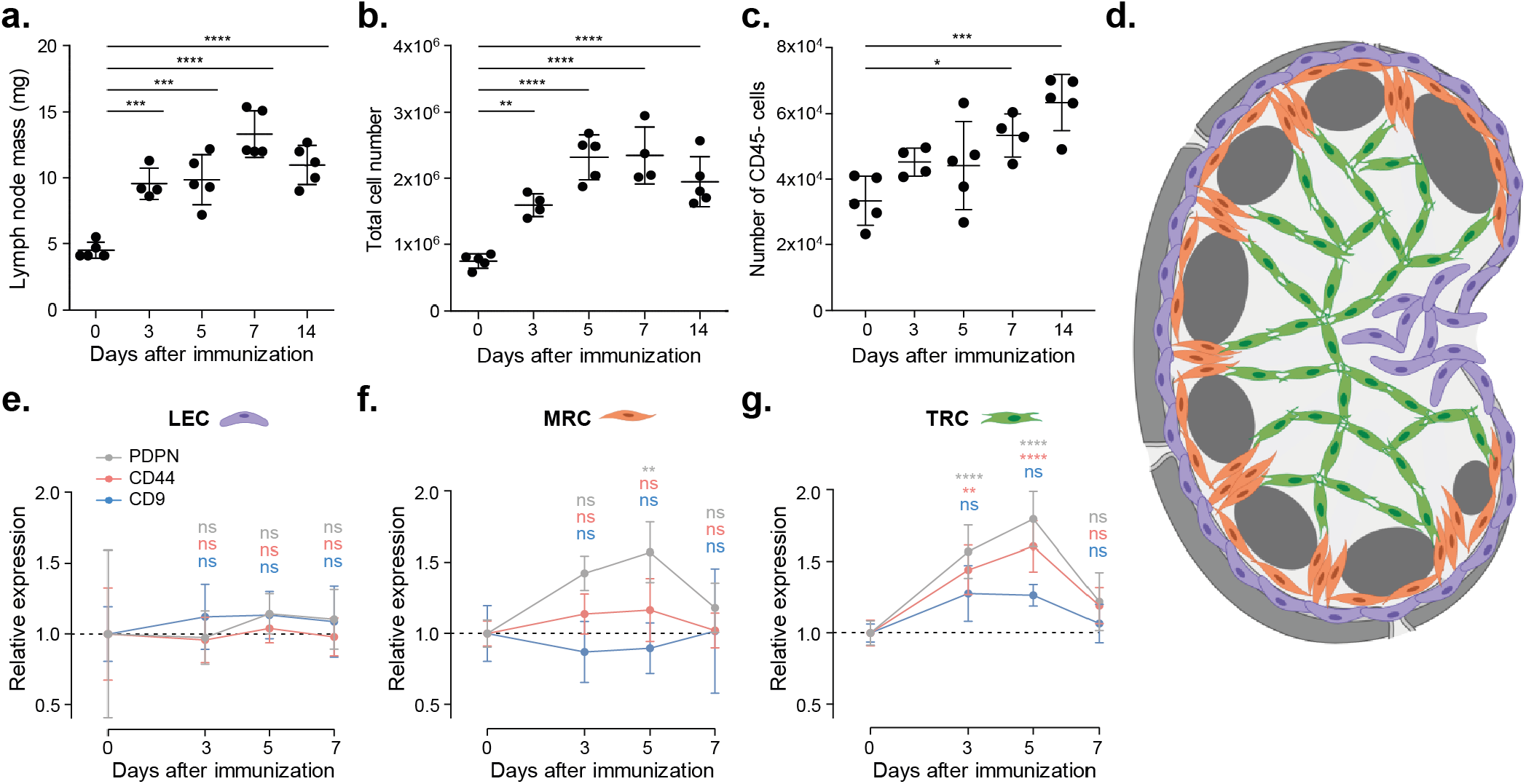
Increased expression of podoplanin and CD44 on TRCs upon *in vivo* immunisation. **a.** Mass (mg) of dissected inguinal lymph nodes. **b-c., e-g.** Analysis by flow cytometry of cell suspensions of inguinal lymph nodes from C57BL/6 mice immunized with IFA/OVA for indicated time points. FRC population is based on total lymph node cell count x (percentage of live cells) x (percentage of CD45^-^ cells). Gating strategy is included in Supplementary Figure 1b. **b.** Total live cell number is based on the live cell gate as measured by flow cytometry. **c.** Number of CD45^-^ cells. **d** Schematic representation of location of different lymph node stromal cell subsets: lymphatic endothelial cell (LEC; purple), marginal zone reticular cell (MRC; orange), and T-cell zone reticular cell (TRC; green). **e-g.** Podoplanin (PDPN; grey), CD44 (pink), and CD9 (blue) surface expression on LECs (CD31^+^podoplanin^+^; **e**), MRCs (CD31^-^podoplanin^+^MAdCAM-1^+^; **f**) and TRCs (CD31^-^podoplanin^+^MAdCAM-1^-^; **g**) as measured by flow cytometry. Protein expression is normalized to day 0. n=4-5 mice per time point. \**p*<0.05, \*\**p*<0.01, \*\*\**p*<0.001, \*\*\*\**p*<0.0001. ns, not significant.

Since podoplanin is induced to cluster in cholesterol-rich regions upon CLEC-2 binding (2), we hypothesised that membrane partner proteins of podoplanin may also be required to downregulate podoplanin-driven contractility for FRC network elongation. A known partner protein of podoplanin, the hyaluronic acid receptor CD44 (19, 20), is also specifically upregulated on TRCs, but not on MRCs or LECs (Fig. 1e-g). Another podoplanin partner protein, tetraspanin CD9 (21), does not significantly change on any of the lymphoid stromal cell subtypes (Fig. 1e-g), but there is a trend towards higher expression on TRCs (Fig. 1g). TRCs are required to elongate upon immune activation, permitting space for rapidly increasing T-cell populations (1, 2). Since podoplanin levels are increased during a phase of lymph node expansion when TRCs are less contractile, we hypothesise that podoplanin may play additional roles on TRCs during an immune response. Interactions with partner proteins CD44 and potentially CD9 may facilitate alternative and unreported podoplanin functions.

### Expression of podoplanin, CD44 and CD9 on FRCs is coregulated by CLEC-2

Investigating the roles of podoplanin, CD44 and CD9 specifically on TRCs *in vivo* is technically challenging since these proteins are broadly expressed in many cell types and carry out essential functions in development and homeostasis. To investigate the functions of podoplanin, CD44 and CD9 in TRCs, we utilised an immortalised FRC cell line (2). We can model the TRC responses during acute lymph node expansion by exposing the FRC cell line to recombinant CLEC-2, or model the more prolonged CLEC-2 exposure from migratory DCs arriving into the lymph node over several days (2, 3) using a CLEC-2-Fc-secreting FRC cell line (5).

Both MRCs and TRCs will contact CLEC-2^+^ migratory DCs entering the lymph node (2, 3, 6, 29), and both upregulate podoplanin expression within this early timeframe (Fig. 1f-g). Using our in *vitro* model system, we show that CLEC-2 binding is sufficient to increase podoplanin protein expression in FRCs (Fig. 2a), suggesting that regulation of podoplanin by CLEC-2 occurs either by inhibiting protein degradation or at the transcriptional level. Indeed, we find in RNA sequencing (RNAseq) data (5) that podoplanin (*Pdpn)* mRNA levels are increased upon short-term CLEC-2 stimulation (Fig. 2b). Furthermore, CLEC-2 stimulation also increases *Cd44* mRNA levels (Fig. 2b), which mimics the increased expression on TRCs during lymph node expansion (Fig. 1g). Podoplanin shRNA knockdown (PDPN KD) FRCs are used as negative control, and we find that both *Cd44* and *Cd9* expression is reduced when podoplanin expression is knocked down. Despite these transcriptional changes, surface expression of podoplanin and CD44 is not increased in the CLEC-2-expressing FRC cell line (Fig. 2c), suggesting that, although more mRNA and protein is produced (Fig. 2a-b), surface expression is already maximal. The RNAseq data show a decrease in *Cd9* mRNA expression upon short-term CLEC-2 stimulation (Fig. 2b). In PDPN KD FRCs, both CD44 and CD9 surface protein levels are reduced (Fig. 2b-c), indicating a degree of co-expression between these partner proteins. We investigated this interdependence by generating CD44 and CD9 knockout (KO) FRC cell lines, and find that knockout of either CD44 or CD9, or both proteins (CD44/CD9 DKO), results in approximately 25% reduction of podoplanin surface expression compared to control FRCs (Fig. 2d). These data suggest that the availability of these two partner proteins impacts podoplanin expression levels at the plasma membrane, providing additional evidence that podoplanin, CD44 and CD9 are coregulated.

**Fig. 2.**
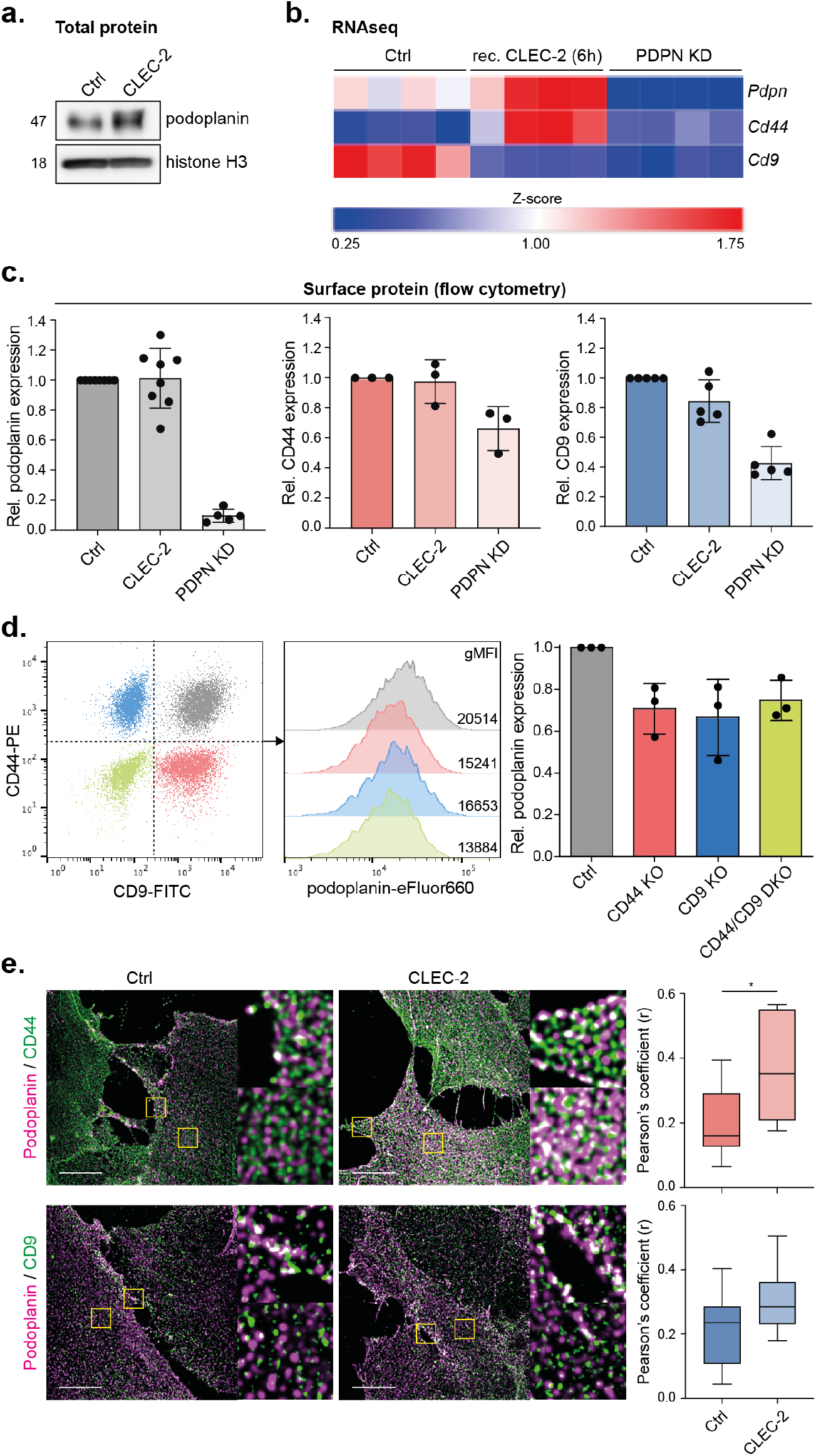
Expression of podoplanin, CD44 and CD9 on FRCs is coregulated by CLEC-2. **a.** Western blot analysis of total podoplanin (PDPN) expression in Ctrl or CLEC-2-Fc-expressing FRC cell lines. Histone H3 is used as loading control. Representative data from n=3 biological replicates is shown. **b.** RNAseq expression (5) of *Pdpn* (top row), *Cd44* (middle row) and Cd9 (bottom row) in unstimulated (left), or short-term stimulated (6h recombinant CLEC-2-Fc) Ctrl FRCs (middle), or podoplanin shRNA knockdown (PDPN KD) FRCs (right). mRNA expression is analysed as Z-score compared to *Pdpn* expression in unstimulated Ctrl FRCs. n=4 biological replicates per cell type or stimulation. **c.** Podoplanin (left), CD44 (middle) and CD9 (right) surface expression on control (Ctrl; dark), CLEC-2-expressing (CLEC-2; lighter), and PDPN KD (light) FRCs by flow cytometry. Protein expression is normalized to the level in Ctrl FRCs for each individual experiment. Data shown as mean+/-SD with dots representing biological replicates (n=3-8). **d.** Podoplanin surface expression on indicated FRCs by flow cytometry. Podoplanin expression is normalized to the level in Ctrl FRCs for each individual experiment. Data shown as mean+/-SD with dots representing biological replicates (n=3). **e.** Left panel: Immunofluorescence of podoplanin (magenta) and CD44 (top; green) or CD9 (bottom; green) in control (Ctrl) or CLEC-2-expressing (CLEC-2) FRCs. Maximum Z stack projections of representative images are shown. The scale bars represent 10 microns. Yellow boxes indicate zoomed in areas. Right panel: Co-localisation of podoplanin with CD44 (red) or CD9 (blue) in Ctrl and CLEC-2 FRCs as measured by Pearson’s coefficient (r). \**p*=0.0476.

It is reported that podoplanin and CD44 can directly interact via either their extracellular domains (19), or their transmembrane and cytoplasmic domains (20). Podoplanin and CD9 are also confirmed as direct partner proteins, interacting through the transmembrane domain of podoplanin and CD9 transmembrane domains 1 and 2 (21). We examined colocalization of podoplanin/CD44 and podoplanin/CD9 complexes on the plasma membrane of FRCs. In steady-state, unstimulated FRCs, both podoplanin/CD44 and podoplanin/CD9 complexes partially colocalise predominantly at the cell periphery (Fig. 2e). CLEC-2-expressing FRCs show significantly increased colocalization of podoplanin/CD44 broadly across the whole cell membrane (Fig. 2e). The abundance of podoplanin/CD9 complexes is not significantly alteredby CLEC-2 stimulation and complexes remain localised to the cell periphery. Thus, upon CLEC-2 stimulation, podoplanin/CD44 and podoplanin/CD9 complexes reside in different subcellular locations, supporting a model of distinct pools of podoplanin on the FRC cell membrane, which may have different functions.

We show that CLEC-2 stimulation of FRCs is sufficient to regulate expression of podoplanin, CD44 and CD9 mRNA and protein, and furthermore regulates their colocalization at the cell membrane. Our data suggest that contact between CLEC-2^+^ DCs and lymphoid fibroblasts would be sufficient to regulate the expression of these surface markers *in vivo* (Fig. 1f-g).

### CD44 and CD9 balance podoplanin-driven FRC contractility

Podoplanin drives actomyosin contractility in FRCs (2, 3). It has been shown that podoplanin-mediated contractility can be counterbalanced by CD44 overexpression (2, 19), which coincides with podoplanin relocalising to cholesterol-rich membrane domains (2). Indeed, cholesterol depletion in FRCs results in hypercontractility and cell rounding in a podoplanin-dependent manner (2). Tetraspanins are predicted to have an intramembrane cholesterol binding pocket controlling their activity (30). We hypothesise that podoplanin activity in FRCs is altered through changing microdomain location in the plasma membrane, mediated by its membrane partner proteins CD44 and CD9. The reported function of podoplanin in FRCs is promoting actomyosin contractility (2, 3). Excess contractility can cause cells to round up, overcoming adhesions, and in the extreme result in membrane blebbing phenotypes (31). CLEC-2 upregulates podoplanin in FRCs (Fig. 2a-b), yet actomyosin contractility is reduced. We tested the capacity for CD44 or CD9 to control podoplanin-driven contractility. In the absence of either CD44 or CD9, FRCs remain spread and exhibit F-actin stress fibres similarly to control cells (Fig. 3a-b), indicating that a balance between contraction, protrusion and adhesion is maintained. However, CD44/CD9 DKO cells round up and contract (40%) (Fig. 3a-b), further quantified by approx. 50% reduction in cell area (Fig. 3c). Hypercontractility in CD44/CD9 DKO FRCs is podoplanin-dependent, since CLEC-2-expressing CD44/CD9 DKO FRCs remain spread (Fig. 3a-c). CD44/CD9 DKO cells express lower podoplanin levels (Fig. 2d) yet contract more than control cells (Fig. 3b), suggesting that both CD44 and CD9 can temper podoplanin-driven contractility. We tested this hypothesis directly by overexpressing PDPN-CFP in combination with CD44 or CD9 GFP-fusion proteins in Ctrl FRCs (Fig. 3d). Over-expression of PDPN-CFP alone induces Ctrl FRCs to round up and exhibit membrane blebs (Fig. 3d). Co-transfecting CD44-GFP or CD9-GFP, and not a GFP control plasmid, rescues this hypercontractile phenotype, and FRCs overexpressing both podoplanin and CD44 or CD9 remain spread (Fig. 3d-f). These data indicate that control of podoplanin-driven FRC contractility requires a balanced expression between podoplanin and its partner proteins CD44 and CD9.

**Fig. 3.**
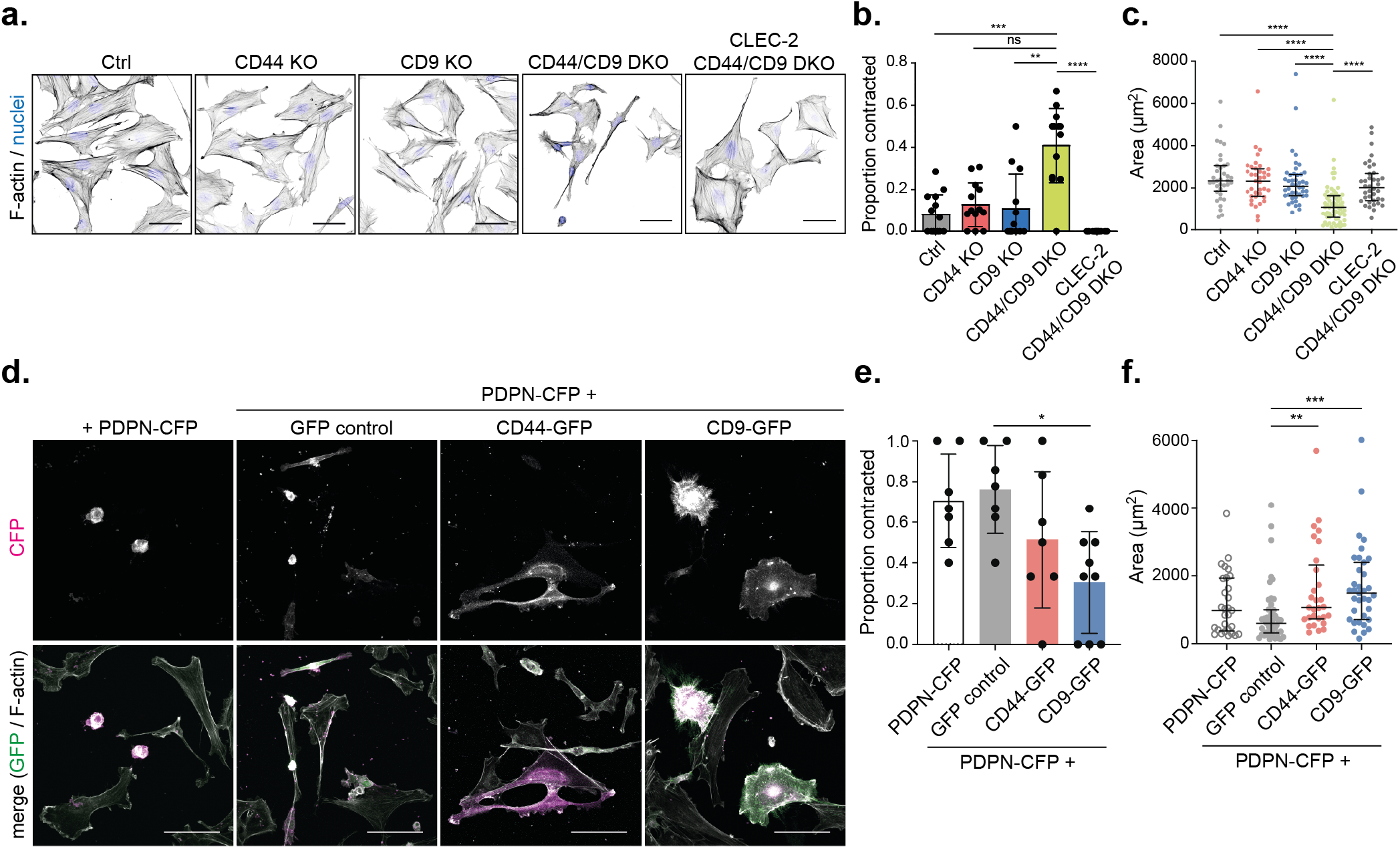
CD44 and CD9 balance podoplanin-mediated FRC hypercontractility. **a.** Immunofluorescence of F-actin (black) and cell nuclei (blue) in indicated FRC cultures. Maximum Z stack projections of representative images are shown. The scale bars represent 50 microns. **b.** Proportion of contracted cells in indicated FRC cultures. Data shown as mean+/-SD with dots representing n=11-15 images per cell line collated from 3-4 biological replicates. \*\**p*=0.0023, \*\*\**p*=0.0009, \*\*\*\**p*<0.0001. ns, not significant (*p*=0.0528). **c.** Area (*μ*m^2^) of indicated FRCs. Data shown as median with interquartile range with dots representing individual cells from 3-4 biological replicates. \*\*\*\**p*<0.0001. **d.** Immunofluorescence of CFP (magenta), GFP (green) and F-actin (white) in Ctrl FRCs transfected with PDPN-CFP alone, or co-transfected with GFP control, CD44-GFP, or CD9-GFP. Maximum Z stack projections of representative images are shown. The scale bars represent 80 microns. **e.** Proportion of contracted cells in Ctrl FRCs transfected with PDPN-CFP alone (white), or co-transfected with GFP control (grey), CD44-GFP (red), or CD9-GFP (blue). Data shown as mean+/-SD with dots representing n=7-9 images per cell line collated from 3-4 biological replicates. \**p*=0.0186. **f.** Area (*μ*m^2^) of Ctrl FRCs transfected with PDPN-CFP alone (white), or co-transfected with GFP control (grey), CD44-GFP (red), or CD9-GFP (blue). Data shown as median with interquartile range with dots representing individual cells from 3 biological replicates.

### FRC motility and polarity are controlled by podoplanin

We show that podoplanin-driven contractility is reduced by the availability of CD44 and CD9 (Fig. 3). However, these data do not explain why podoplanin expression is upregulated in TRCs during acute lymph node expansion (Fig. 1), a phase of tissue remodelling when the fibroblastic reticular network is elongating and is less contractile (2, 3). We asked whether podoplanin controls additional aspects of fibroblastic reticular network function, beyond contractility. To maintain network integrity during acute lymph node expansion, FRCs elongate, reduce adhesion and detach from the underlying conduit network (5). To accommodate the expanding lymphocytes, the FRC network must be flexible and motile, yet retain network connectivity (2, 3, 5). We tested the requirement for podoplanin, CD44 and CD9 in FRC motility and polarity.

To model FRC motility, we studied the displacement of FRCs in 2D time-lapse assays. It is notable that even as single cells, *in vitro*, control FRCs were not highly migratory, moving at speeds <0.2 /min (Fig. 4a-b), which is consistent with their ability to preserve network integrity *in vivo*. PDPN KO FRCs show increased motility compared to control FRCs, measured by both cumulative distance (Fig. 4a) and displacement (Fig. 4b), which is unaffected by additional knock-out of CD44 (PDPN/CD44 DKO) or CD9 (PDPN/CD9 DKO) (Fig. 4a-b). However, in podoplanin^+^ FRCs, both CD44 KO and CD9 KO increase motility in a non-redundant manner (Fig. 4a-b). These data suggest that podoplanin, CD44 and CD9 may have roles to play in maintaining the non-migratory phenotype of FRCs.

**Fig. 4.**
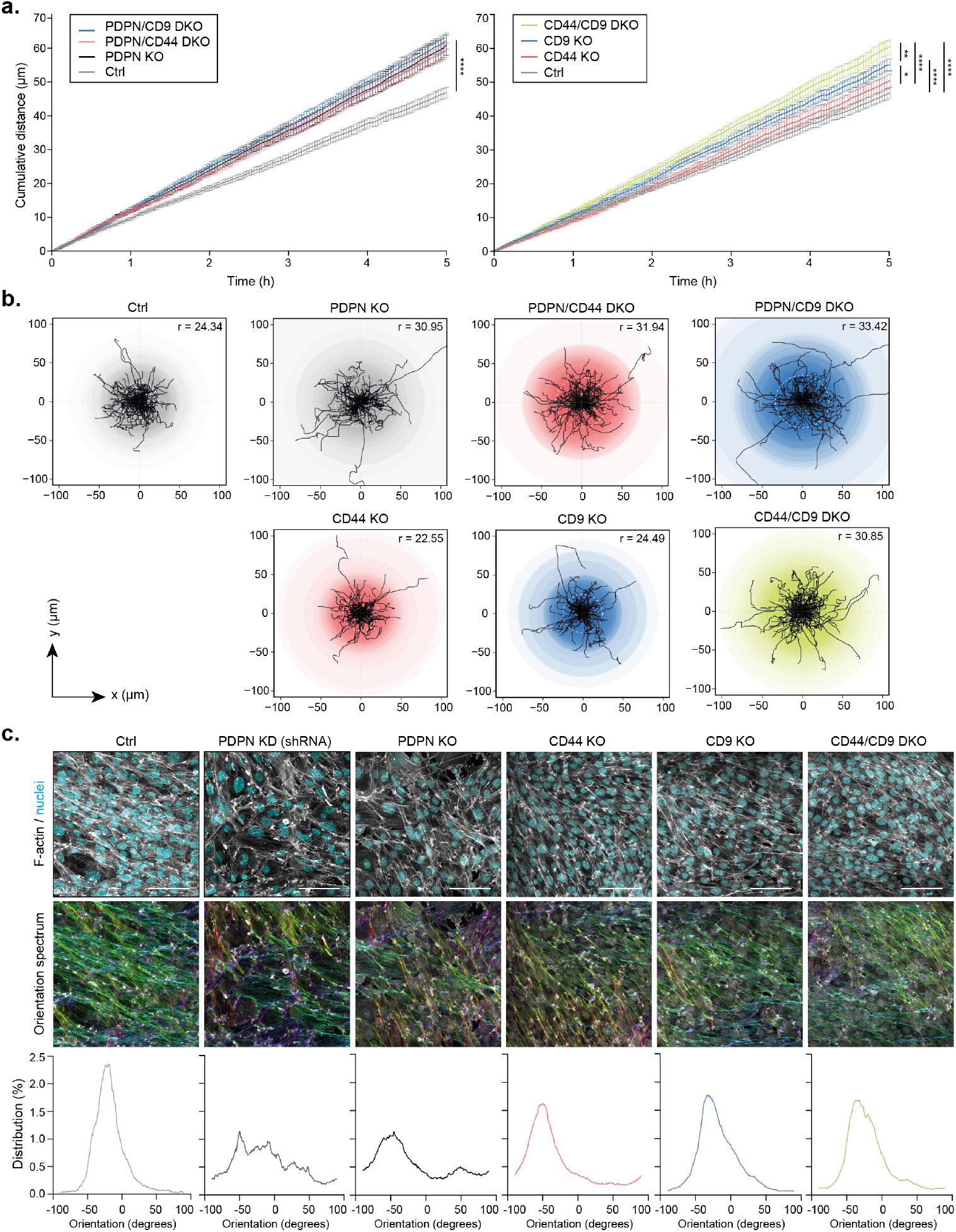
Podoplanin also controls FRC motility and polarity. **a.** Mean cumulative distance (in *μm)* travelled by FRCs as indicated in the legend (top left of each graph) over the course of 5 hours. Data shown as mean+/-SEM for each timepoint representing n = 71-118 cells per cell line. \**p*<0.05, \*\**p*<0.01, \*\*\*\**p*<0.0001. **b.** Individual tracks of FRCs during the first 120 minutes of the data presented in **a.** Semi-transparent coloured circles indicate the maximum distance from the origin reached by each cell. The median maximum distance per condition is depicted in the top right corner and indicated as a dotted white circle in each plot. **c.** Top: Indicated FRCs were cultured until full confluency was reached, and stained for F-actin (grey) and cell nuclei (cyan). Maximum Z stack projections of representative images are shown. The scale bars represent 100 microns. Middle: FRC-FRC alignment analysed using OrientationJ plugin based on F-actin staining. The colours in the Orientation spectrum indicate the 2D orientation (−180 to 180 degrees) per pixel. Bottom: Histograms showing the distribution (in percentage) of the pixel orientation (in degrees) per image.

Unlike other fibroblast populations which exhibit contact inhibition of locomotion (CIL) (32, 33), repolarising and migrating away from neighbouring cells upon contact, FRCs physically connect to form an intricate multi-cellular network (2, 34–36). Network connectivity is maintained and prioritised throughout the early phases of lymph node expansion (2, 3, 34). It is unknown how FRCs overcome CIL to form stable connections with their neighbours. Our data show that control podoplanin^+^ FRCs align with each other in *in vitro* cultures, however, both PDPN KD and PDPN KO FRCs lack this alignment (Fig. 4c). Neither CD44 KO, CD9 KO, or CD44/CD9 DKO FRCs alter FRC alignment (Fig. 4c). Together, these data indicate that podoplanin inhibits FRC motility and promotes alignment, showing for the first time that podoplanin plays an important role in FRC function beyond actomyosin contractility.

### CD9 and CD44 modulate FRC-FRC interactions

Next, we wanted to investigate FRC-FRC interactions in more detail. We noticed a 25% increase in cumulative displacement in CD9 KO FRCs compared to control and CD44 KO FRCs (Fig. 4a), suggesting an independent role for CD9 on FRC motility. In other cell types, CD9 is known to be important for cell migration and adhesion, and cell fusion (24–26). Therefore, we further investigated the role for CD9 in FRC motility and FRC-FRC interactions.

We compared Arp2/3^+^ protrusions, as a readout of actively protruding plasma membrane (37), in CD44 KO, CD9 KO and PDPN KO FRC cell lines . FRCs lacking podoplanin, CD44, or CD9 show increased membrane localisation of ARPC2, part of the Arp2/3 complex, compared to control FRCs (Fig. 5a), which is in line with increased motility observed in each of these cell lines (Fig. 4a-b). CD9 KO FRCs have broad ARPC2^+^ protrusions, covering most of the plasma membrane (Fig. 5a). This increased protrusion is also observed in PDPN/CD9 DKO cells, but not in PDPN KO cells (Fig. 5a), indicating that this phenotype is CD9-dependent, and independent of podoplanin function. Control FRCs meet and interact with their neighbouring FRCs forming connections between one another with little overlap of membranes (Fig. 5a-c). However, CD9 KO and PDPN/CD9 DKO FRCs continued to protrude and spread, as such overlapping neighbouring cells (Fig. 5a-c). In contrast, CD44 KO FRCs showed reduced areas of overlap between neighbouring cells (Fig. 5a-c). Our data indicate that CD9 and CD44 play opposing roles to modulate how FRCs make contacts with neighbouring cells, and we suggest that CD9 and CD44 expression by FRCs may facilitate FRC network formation in vivo.

**Fig. 5.**
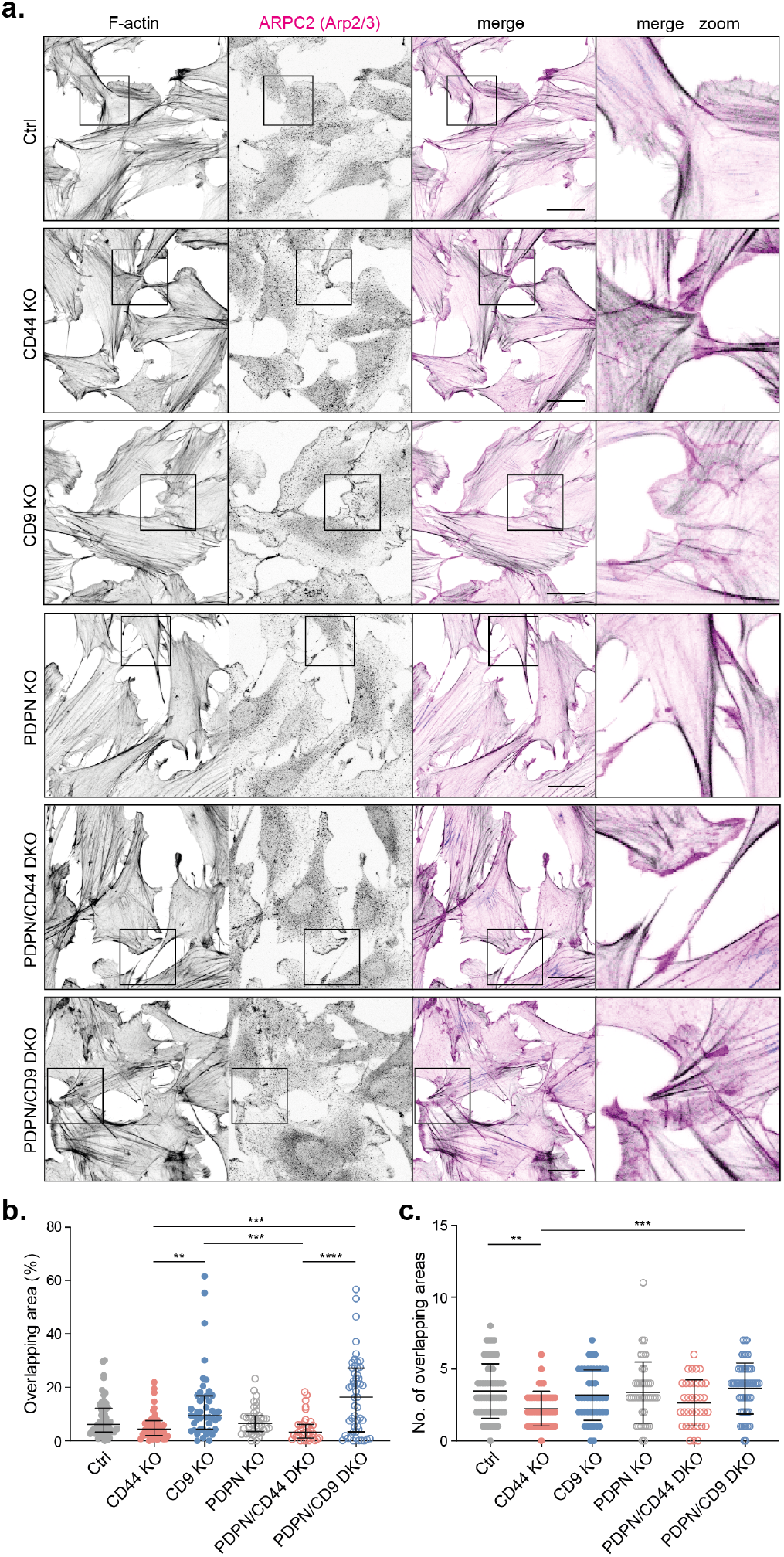
CD9 and CD44 regulate FRC-FRC interactions. **a.** Immunofluorescence of F-actin (black) and Arp2/3^+^ protrusions (visualized by staining the ARPC2 subunit; magenta) in indicated FRC cultures. Maximum Z stack projections of representative images from n=2 biological replicates are shown. The scale bars represent 30 microns. **b-c.** Percentage of total overlapping area (**b**) and number of overlapping areas (**c**) for indicated FRCs. Dots represent single FRCs. n=36-57 cells in total from 2 biological replicates. Error bars represent median with interquartile range. \*\**p*<0.01, \*\*\**p*<0.001, \*\*\*\**p*<0.0001.

### CD44 and CD9 control podoplanin-dependent FRC responses to CLEC-2^+^ DCs

Podoplanin was first described as a ligand promoting both platelet aggregation and DC migration (6, 13). We next tested whether CD44 or CD9 expression by FRCs is required for podoplanin ligand function using 3D co-culture of FRCs with bone marrow-derived CLEC-2^+^ DCs. Contact with podoplanin^+^ FRCs induces DCs to extend protrusions, in a CLEC-2 (6) and tetraspanin CD37-dependent manner (38). DCs co-cultured with PDPN KD FRCs do not spread or make protrusions (Fig. 6a). However, co-culture of DCs with CD44 KO or CD9 KO FRCs does not hamper DC responses (Fig. 6a). Furthermore, the increase in morphology index (perimeter^2^/4*π* area) is equivalent to DCs co-cultured with control FRCs (Fig. 6b). As such, podoplanin ligand function is not dependent on CD44 or CD9 expression on FRCs. This is in agreement with published data showing that soluble recombinant podoplanin-Fc can induce DC protrusions (6, 38).

**Fig. 6.**
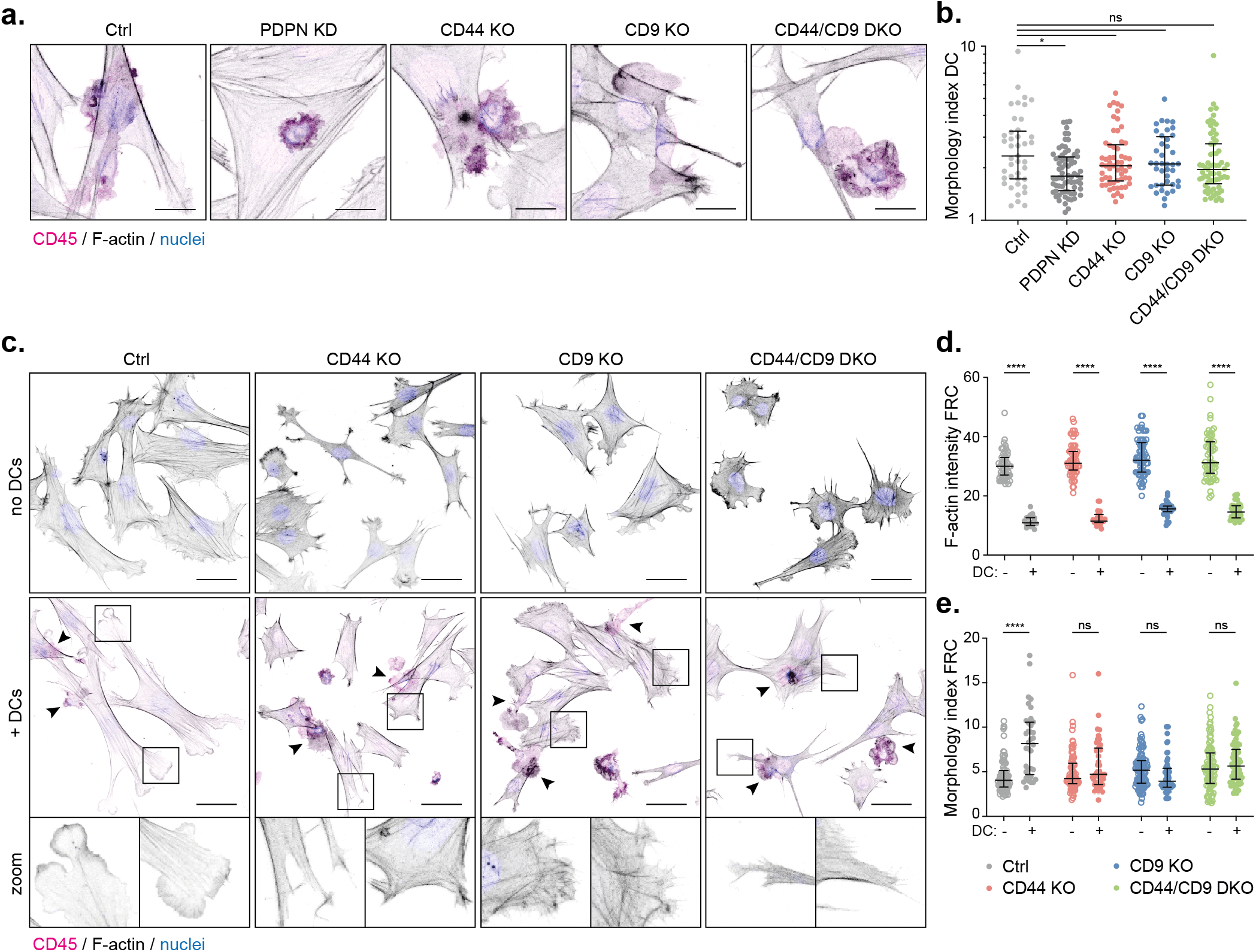
Podoplanin, CD44 and CD9 are required for FRCs to respond to CLEC-2^+^ DCs. **a.** Immunofluorescence of 3D cultures of indicated FRCs with LPS-stimulated bone marrow-derived DCs (CD45^+^; magenta). Maximum Z stack projections of representative images from n=2 biological replicates are shown. The scale bars represent 20 microns. **b.** Morphology index (=perimeter^2^/4*π* area) of DCs in interaction with an FRC. Dots represent single DCs. n=40-72 DCs collated from 2 biological replicates. Error bars represent median with interquartile range. \**p*=0.0149. ns, not significant. Y-axis, log 10 scale. **c.** Immunofluorescence of 3D cultures of FRCs without (upper row) or with (middle and bottom rows) LPS-stimulated bone marrow-derived DCs (magenta). Maximum Z stack projections of representative images from n=2 biological replicates are shown. The scale bars represent 50 microns. **d-e.** F-actin intensity (mean gray value of phalloidin-TRITC staining; **d**), and morphology index (=perimeter^2^/4*π* area; **e**) of indicated FRCs alone (open circles) or in interaction with a DC (closed circles). Dots represent single FRCs. n=36-104 FRCs collated from 2 biological replicates. Error bars represent median with interquartile range. \*\*\*\**p*<0.0001. ns, not significant.

In the lymph node, the fibroblastic reticular network is primed to remodel and elongate during immune responses by contact with CLEC-2^+^ DCs (2, 3). It is known that podoplanin expression by FRCs is required for this signalling pathway, which primarily inhibits actomyosin contractility (2, 3). Since we have shown that CD44 and CD9 both control FRC contractility in a podoplanin-dependent manner (Fig. 3), and may also play roles in FRC motility and FRC-FRC interactions (Fig. 4–5), we ask if CD44 and/or CD9 expression is required by FRCs to respond to CLEC-2^+^ DCs. Binding of CLEC-2^+^ DCs to FRCs drives elongation and induction of multiple protrusions, which *in vivo*, is required for acute lymph node expansion during adaptive immune responses (2, 3). FRCs respond to CLEC-2^+^ DCs *in vitro* by forming lamellipodia-like actin-rich protrusions in multiple directions (Fig. 6c), and by a reduction in F-actin fibres (Fig. 6c-d). We interpret this response as reduced actomyosin contractility, and a concurrent increase in actin polymerisation driving protrusions. This is quantified by increased morphology index (perimeter^2^/4*π* area; Fig. 6e). Strikingly, both CD44 KO and CD9 KO FRCs fail to form lamellipodia in response to DC contact (Fig. 6c). CD44 KO FRCs exhibit small protrusions with F-actin ‘spikes’, whereas CD9 KO FRCs attempt broader protrusions, but fail to accumulate F-actin at the leading edge (Fig. 6c). These defects are quantified by the lack of increased morphology index in response to CLEC-2^+^ DCs (Fig. 6e). However, we still observe a DC-induced reduction in F-actin fibres in CD44 KO and CD9 KO FRCs, as well as in CD44/CD9 DKO FRCs (Fig. 6c-d), confirming that CLEC-2^+^ DCs still contact the FRCs and can inhibit contractility pathways (Fig. 6a). We conclude that both CD44 and CD9 participate in podoplanin-dependent spreading. Indeed, even before contact with DCs, CD44 KO and CD9 KO FRCs are spread over a smaller area compared to control FRCs (Fig. 6c), suggesting that CD44 and CD9 also act to balance podoplanin-driven contractility and protrusion formation in steady state. These data lead us to conclude that the induction of spreading and the formation of lamellipodia protrusions in response to DC contact is an active podoplanin-dependent process, which requires both CD44 and CD9 in a non-redundant fashion.

Overexpression of podoplanin drives hypercontractility (2, 3), yet when FRCs are spreading and elongating during the initiation of lymph node expansion, podoplanin expression is increased (Fig.1). Our data demonstrate that podoplanin is involved in multiple FRC functions beyond actomyosin contractility, and that these functions are controlled by podoplanin partner proteins CD44 and CD9.

## Discussion

Lymph node expansion is a transient and reversible process, a cycle of controlled tissue remodelling through each adaptive immune response (4). FRCs shape lymph node architecture (4) and control the balance between actomyosin contractility and spreading/elongation to determine both lymph node structure and size (2, 3, 34). Upon initiation of an adaptive immune response, CLEC-2^+^ migratory DCs inhibit podoplanin-driven FRC contractility to permit lymph node expansion (2, 3). We now show that podoplanin and its partner proteins CD44 and CD9 play key roles in balancing different FRC functions in steady-state, and in response to CLEC-2^+^ DCs.

We show that podoplanin expression by FRCs controls functions beyond contractility. During an immune response, proliferating T cells provide a mechanical strain for TRCs (1), and TRC elongation and spreading is required to preserve network connectivity and stromal architecture, in advance of their proliferation (2, 3). We suggest that podoplanin expression by TRCs is pivotal to their adaptable phenotype. Our data show that podoplanin surface expression increases on TRCs in the first 3-5 days after immunization, a period when contractility through the network is reduced. Furthermore, we find that CLEC-2 stimulation of FRCs upregulates podoplanin mRNA and protein expression. Surface expression of CD44 or CD9, or stimulation with CLEC-2 is able to counterbalance podoplanin-driven contractility. We find that in addition to contractility, podoplanin is required to inhibit lymphoid fibroblast motility and preserve polarity, which would be essential for maintaining FRC network integrity in homeostasis and during lymph node expansion.

In this study, we sought to understand the function of other FRC markers in the normal physiology of immune responses. We find a role for tetraspanin CD9 in controlling FRC-FRC interactions. Tetraspanins are membrane-organizing proteins controlling a variety of cellular processes, including cell-cell interactions, cell migration, and signalling events (22, 23). FRCs lacking CD9 do not detect neighbouring cells, and spread and grow over each other. In other non-lymphoid tissues, CD9 controls cell migration and adhesion, and is required for cell-cell fusion (24–26). We hypothesize that CD9 on FRCs contributes to the formation and preservation of the FRC network. We also found that CD44 surface expression increases on TRCs during the initial phase of lymph node expansion. Furthermore, CLEC-2 stimulation increases colocalisation of podoplanin and CD44 on FRCs. We find that both CD44 and CD9 on FRCs are able to supress podoplanindriven contractility, facilitating alternative FRC phenotypes.

It is previously reported that podoplanin/CLEC-2 binding is necessary for the lymph node to expand in the early phases of an adaptive immune response (2, 3). Importantly, we now show that podoplanin expression by FRCs is not sufficient to respond to CLEC-2^+^ DCs. FRCs in contact with CLEC-2^+^ DCs respond by reducing F-actin cables, making protrusions in multiple directions and spreading over a larger area. It has been assumed that FRC spreading is an indirect event in response to inhibition of actomyosin contractility (2, 3). We find that FRCs lacking either CD44 or CD9 expression still reduce F-actin cables in response to DCs, however, they fail to make protrusions and spread. Our data show that CD44 is involved in broad lamellipodial protrusions, and CD9 regulates filopodial-like protrusions. We conclude that both CD44 and CD9 are required for DC-mediated FRC elongation and spreading, and we now identify loss of actomyosin contractility and spreading as two differently regulated, but linked active processes.

The FRC response to CLEC-2^+^ DCs is multi-faceted, and we need to understand more about CLEC-2-driven podoplanin signalling controlling these various functions. We know that both podoplanin and CD44 bind ERM proteins, which couple membrane proteins to the actin cytoskeleton, driving contractility (2, 3, 7, 27). CLEC-2 binding results in ERM dephosphorylation and decoupling from podoplanin (2, 3). We show that CLEC-2 stimulation results in increased co-localisation of podoplanin and CD44 on FRCs. It is currently unknown if CLEC-2-mediated clustering of podoplanin and CD44 forces uncoupling of ERM proteins, or if dephosphorylation and uncoupling of ERM proteins by an unknown mechanism provides space for clustering of podoplanin/CD44 complexes. Moreover, further studies are required to identify the specific cascade of signalling events downstream the CLEC-2/podoplanin axis facilitating FRC spreading and elongation. Our data suggest that the function of podoplanin and downstream signalling would be determined not only by podoplanin expression level or subcellular localisation, but by the availability of its membrane partner proteins CD44 and CD9. Podoplanin, CD44 and CD9 are broadly expressed and their expression can alter in various pathologies (8, 26, 39). Our findings may further provide insights as to how the balance of their expression may drive cellular functions in pathological conditions.

This study provides a molecular understanding into how FRC functions are controlled in homeostasis and during lymph node expansion. Interactions between podoplanin and two of its known partner proteins, CD44 and CD9 shape dynamic FRC responses, supporting a model of distinct functional protein domains on the FRC plasma membrane. We hypothesize that during homeostasis, lymph node size remains stable via tonic CLEC-2 signalling provided by resident and migratory DCs, maintaining the balance between contraction and protrusion through the FRC network. As the lymph node expands, this mechanical balance in the TRC network is transiently shifted towards cell elongation and protrusion by an influx of CLEC-2^+^ migratory DCs. Our data indicate that TRCs require expression of CD44 and CD9, in addition to podoplanin, to facilitate this shift.

## Materials and methods

Biological materials generated for this study are available upon request to the corresponding author with an MTA where appropriate.

### Mice

Wild-type C57BL/6J mice were purchased from Charles River Laboratories. Female mice were used for *in vivo* experiments and were aged 6-10 weeks. Mice were age matched and housed in specific pathogen-free conditions. All animal experiments were reviewed and approved by the Animal and Ethical Review Board (AWERB) within University College London and approved by the UK Home Office in accordance with the Animals (Scientific Procedures) Act 1986 and the ARRIVE guidelines.

### *In vivo* immunizations

Mice were immunized via subcutaneous injection in the right flank with 100 *μl* of an emulsion of OVA in incomplete (IFA) Freund’s adjuvant (100 *μg* OVA per mouse; Hooke Laboratories). Draining inguinal lymph nodes were taken for analysis by flow cytometry. Lymph nodes were digested as previously described (40), and cells were counted using Precision Count Beads as per supplier’s instructions (Biolegend), and stained for analysis by flow cytometry.

### Cell culture

Control and PDPN KD FRC cell lines (2), and CLEC-2-Fc expressing FRCs (5) are previously described. PDPN KO FRC cell line was generated using CRISPR/Cas9 editing. Control FRC cell line (2) was transfected with pRP[CRISPR]-hCas9-U6>PDPN gRNA 1 plasmid (constructed and packaged by Vectorbuilder; vector ID: VB160517-1061kpr) using Lipofectamine 2000 Transfection Reagent (Thermo Fisher Scientific). We performed three rounds of transfection, and subsequently performed magnetic cell sorting (MACS) using MACS LD columns (Miltenyi Biotec), anti-mouse podoplanin-biotin antibody (clone 8.1.1, eBioscience, 13-5381-82), and anti-biotin microbeads (Miltenyi Biotec, 130-090-485) as per supplier’s instructions to sort PDPN KO FRCs by negative selection. Complete knockout of podoplanin expression was confirmed using quantitative RT-PCR, flow cytometry and Western blot.

CD44 KO, CD9 KO, and CD44/CD9 DKO FRCs were generated using CRISPR/Cas9 editing. Control, CLEC-2-expressing or PDPN KO FRCs were transfected using Attractene Transfection Reagent (Qiagen) with one or both of the following plasmids: pRP[CRISPR]-hCas9-U6>CD44-T3 exon 2 (constructed and packaged by Vectorbuilder; vector ID: VB180119-1369pus), or pRP[CRISPR]-hCas9-U6>CD9 exon 1 (constructed and packaged by Vectorbuilder; vector ID: VB180119-1305adb). Subsequently, CD44 KO, CD9 KO, and CD44/CD9 DKO FRCs underwent two or three rounds of FACS to obtain a full CD44 and/or CD9 KO FRC cell line, which was confirmed using quantitative RT-PCR, flow cytometry and Western blot.

FRC cell lines were cultured in high-glucose DMEM with GlutaMAX supplement (Gibco, via Thermo Fisher Scientific) supplemented with 10% fetal bovine serum (FBS; Sigma-Aldrich), 1% penicillin-streptomycin (P/S) and 1% insulin-transferrin-selenium (both Gibco, via Thermo Fisher Scientific) at 37°C, 10% CO_2_, and passaged using cell dissociation buffer (Gibco, via Thermo Fisher Scientific).

Bone marrow-derived dendritic cells (BMDCs) were generated by culturing murine bone marrow cell suspensions in RPMI 1640 medium (Gibco, via Thermo Fisher Scientific) supplemented with 10% FBS, 1% P/S and 50 *μ*M 2-mercaptoethanol (Gibco, via Thermo Fisher Scientific), and 20 ng/ml recombinant murine granulocyte-macrophage colony-stimulating factor (mGM-CSF, Peprotech, 315-03), as adapted from previously described protocols (41), at 37°C, 5% CO_2_. On day 6, BMDCs were additionally stimulated with 10 ng/ml lipopolysaccharides from *E.coli* (LPS; Sigma-Aldrich, L4391-1MG) for 24 hours.

### DC-FRC co-cultures

FRCs (0.7×10^4^ cells per well) were seeded on 24-well glass-bottomed cell culture plates (Mat-Tek) at 37°C, 10% CO_2_. After 24 hours, LPS-stimulated BMDCs (2.5×10^5^ cells per well) were seeded into 3D collagen (type I, rat tail)/Matrigel matrix (both from Corning, via Thermo Fisher Scientific) supplemented with 10% minimum essential medium alpha medium (MEMalpha, Invitrogen, via Thermo Fisher Scientific) and 10% FCS (Greiner Bio-One) on top of the FRCs. Co-cultures were incubated overnight at 37°C, 10% CO_2_. The next day, co-cultures were fixed and stained for analysis by microscopy.

### Flow cytometry

For analysis of lymph nodes from *in vivo* immunisations by flow cytometry, 3×10^6^ cells were incubated with purified rat IgG2b anti-mouse CD16/32 receptor antibody as per supplier’s instructions (Mouse BD Fc-block, clone 2.4G2, BD Biosciences, 553141) for 20 minutes at 4°C. Cells were stained with the following primary mouse antibodies (1:100 dilution) for 30 minutes at 4°C: CD45-BV750 (clone 30-F11, Biolegend, 103157), CD31-PE-Cy5.5 (clone MEC 13.3, BD Biosciences, 562861), podoplanin-PE (clone 8.1.1, BD Biosciences, 566390), CD44-BV605 (clone IM7, BD Biosciences, 563058), CD9-FITC (clone MZ3, Biolegend, 124808) and MAdCAM-1-BV421 (clone MECA-367, BD Biosciences, 742812). Cells were washed with PBS and stained with Zombie Aqua fixable live-dead kit as per supplier’s instructions (Biolegend, 423101) for 30 minutes at 4°C. Next, cells were fixed using Biolegend Fixation/Permeabilization Buffer as per supplier’s instructions (Biolegend, 421403). Samples were analysed on BD Symphony A5 equipped with 355 nm, 405 nm, 488 nm, 561 nm and 638 nm lasers. Acquisition was set to 5×10^5^ single, live CD45^+^ cells. FRC cell number was based on their percentage within the CD45^-^ cells.

Single-cell suspensions of FRC cell lines were incubated with FcR blocking reagent (Miltenyi Biotec) as per supplier’s instructions, followed by 30 minutes staining on ice with the following primary mouse antibodies diluted in PBS supplemented with 0.5% BSA and 5mM EDTA: hamster anti-podoplanin-eFluor660 (clone 8.1.1, 1:200, eBioscience, 50-5381-82), rat anti-CD44-PE (clone IM7, 1:50, BD Biosciences, 553134), and/or rat anti-CD9-FITC (clone MZ3, 1:50, Biolegend, 124808). Stained cells were analyzed using FACSDiva software and LSR II flow cytometer (both BD Biosciences). All flow cytometry data was analysed using FlowJo Software version 10 (BD Biosciences).

### Western blot

Ctrl or CLEC-2-Fc FRCs were plated in a 6-well culture plate (1×10^5^ cells per well). After 24 hours, the culture plate was placed on ice and cells were washed twice with cold PBS. Cells were lysed in 100 *μl* 4x Laemmli lysis buffer (Bio-Rad) and collected using cell scraper. Samples were separated by reducing 10% SDS-polyacrylamide gel electrophoresis. Western blots were incubated with rat antimouse podoplanin (clone 8F11, 1:1000, Acris Antibodies, AM26513AF-N), or mouse anti-Histone H3 (1:2000, Abcam, ab24824) as loading control, in PBS supplemented with 1% skim milk powder and 0.2% BSA overnight at 4°C, followed by staining with appropriate HRP-conjugated secondary antibodies (Abcam) for 2 hours at RT. Western blots were developed using Luminata Crescendo Western HRP substrate (Merck Millipore) and imaged on ImageQuant LAS 4000 mini (GE Healthcare Life Sciences).

### Transient transfection

FRCs were plated on glass coverslips in a 6-well culture plate (2.5×10^4^ cells per well) one day before transfection with PDPN-CFP and CD44-GFP (2), CD9-GFP (kind gift from Dr. Sjoerd van Deventer and Prof. Annemiek van Spriel, Radboudumc, Nijmegen, NL) or GFP control plasmid (0.5*μ*g DNA per plasmid) using Attractene transfection reagent (Qiagen) as per supplier’s instructions. 24 hours post-transfection, FRCs were fixed for analysis of cell contractility by microscopy.

### Immunofluorescence

FRCs were seeded on glass coverslips for 24 hours at 37°C, 10% CO_2_. Next, cells were fixed in 3.6% formaldehyde (Sigma-Aldrich; diluted in PBS), and subsequently blocked in 2% BSA in PBS and stained for 1 hour at RT with the following primary mouse antibodies: hamster anti-podoplanin-eFluor660 (clone 8.1.1, 1:200, eBioscience, 50-5381-82), rat anti-CD44 (clone IM7, 1:200, BD Biosciences, 553131), rat anti-CD9-eFluor450 (clone KMC8, 1:200, eBioscience, 14-0091-82), or rabbit anti-p34-Arc/ARPC2 (Arp2/3, 1:100, Merck, 07-227). This was followed by incubation with appropriate Alexa Fluor-conjugated secondary antibodies (1:500, Invitrogen, via Thermo Fisher Scientific) for 1 hour at RT. F-actin and cell nuclei were visualized using respectively phalloidin-TRITC (P1951-1MG) and DAPI (D9542-1MG; both 1:500 dilution, both from Sigma-Aldrich) incubated for 15 minutes at RT, and coverslips were mounted in Mowiol (Sigma-Aldrich). Cells were imaged on a Leica SP5 or SP8 confocal microscope using respectively HCX PL APO or HC PL APO CS2 /1.4 63x oil lenses.

DC-FRC cultures were fixed with AntigenFix (DiaPath, via Solmedia) for 3 hours at room temperature (RT), followed by permeabilization and blocking with 2% bovine serum albumin (BSA; Sigma-Aldrich) and 0.2% Triton X-100 in phosphate-buffered saline (PBS) for 2 hours at RT. Subsequently, BMDCs were stained using rat anti-mouse CD45-AF647 (clone 30-F11, 1:250, Biolegend, 103123), and Factin and cell nuclei were visualized using respectively phalloidin-TRITC (P1951-1MG) and DAPI (D9542-1MG; both 1:500 dilution, both from Sigma-Aldrich). Co-cultures were imaged on a Leica SP5 confocal microscope using HCX PL APO /1.25 40x oil lenses.

Images were analysed using Fiji/ImageJ software. Z stacks (0.5 *μ*m/step) were projected with ImageJ Z Project (maximum projection). Podoplanin/CD44 and podoplanin/CD9 co-localisation was analysed using JACoP plugin (42). Proportion of contracted cells was quantified by analysing the number of contracted/blebbing cells (based on F-actin staining) compared to total number of cells per field of view. Cell area of FRCs was analysed by manually drawing around the cell shape using F-actin staining. FRC alignment was analysed using OrientationJ plugin (43). Overlapping cell area was determined based on ARPC2 and F-actin staining to determine cell periphery and separate cells, respectively. Morphology index (=perimeter^2^/4*π* area) was calculated using the area and perimeter of BMDCs or FRCs by manually drawing around the cell shape using F-actin staining.

### Live imaging and analysis

Control and PDPN, CD44 and/or CD9 KO FRC cells (5×10^4^ cells per well) were seeded in 12-well plates and incubated overnight at 37°C, 10% CO_2_. The next day, cells were imaged every 10 minutes on a Nikon Ti inverted microscope fitted with a Nikon DS-Qi2 CMOS camera and a controlled 37°C and 5% CO_2_ atmosphere chamber. For motility analysis, cells were manually tracked using Fiji/ImageJ Manual Tracker plugin. Individual cells were tracked by clicking the centre of the nucleus from the start of a file or immediately after cell division, and until the next division, the end of the file or until the nucleus reached the boundaries of the image field. The distance travelled between frames and position per frame was calculated by Fiji. R was used to calculate the cumulative distance travelled per cell and to plot the individual trajectories (ref: R Core Team (2020). R: A language and environment for statistical computing. R Foundation for Statistical Computing, Vienna, Austria. https://www.R-project.org/).

### Statistics

Statistical differences between two groups were determined using unpaired Student’s t-test (one-tailed), or, in the case of non-Gaussian distribution, Mann-Whitney test. Statistical differences between two different parameters were determined using one-way ANOVA with Tukey’s multiple comparisons test. Statistical differences between more than two groups were determined using two-way ANOVA with Tukey’s multiple comparisons test, or, in the case of non-Gaussian distribution, Kruskal-Wallis test with Dunn’s multiple comparisons. Statistical tests were performed using GraphPad Prism software (version 7), and differences were considered to be statistically significant at *p* ≤ 0.05.

### Data availability

RNAseq data (5) (used in Fig. 2b) are publicly available through UCL research data repository: 10.5522/04/c.4696979.

## ACKNOWLEDGEMENTS

We thank Prof. Erik Sahai (The Francis Crick Institute, UK), Prof. Annemiek van Spriel (RIMLS, Radboudumc, NL), Dr. Louise Cramer (MRC-LMCB, UCL, UK), Dr. Christopher Tape (UCL Cancer Institute, UK), and Harry Horsnell (PhD student at MRC-LMCB, UCL, UK) for critical reading of the manuscript. We thank the core staff at the UCL Cancer Institute Flow Cytometry Facility for sorting CRISPR/Cas9 edited FRC cell lines and for use of BD Symphony A5 flow cytometer. This work is supported by a Dutch Research Foundation (NWO) Rubicon Postdoctoral Fellowship (019.162LW.004; to C.M.d.W), European Research Council Starting Grant (LNEXPANDS; to S.E.A), Cancer Research UK Career Development Fellowship (CRUK-A19763; to S.E.A) and Medical Research Council (MC-U12266B).

## AUTHOR CONTRIBUTIONS

C.M.d.W. and S.E.A. designed the study and wrote the manuscript; S.M. performed in vivo protein expression analysis; L.M. investigated changes in and transcriptional control of podoplanin expression upon CLEC-2; J.C.R. analysed overexpression and live imaging data; A.C.B. generated CLEC-2-Fc and (inducible) PDPN-KO FRC cell lines and assisted with cell sorting; V.G.M. performed FRC transcriptome analysis; C.M.d.W. generated CD44 KO, CD9 KO, and CD44/CD9 DKO FRC cell lines, and performed and analysed all other experiments.

---

**Fig. S1.**
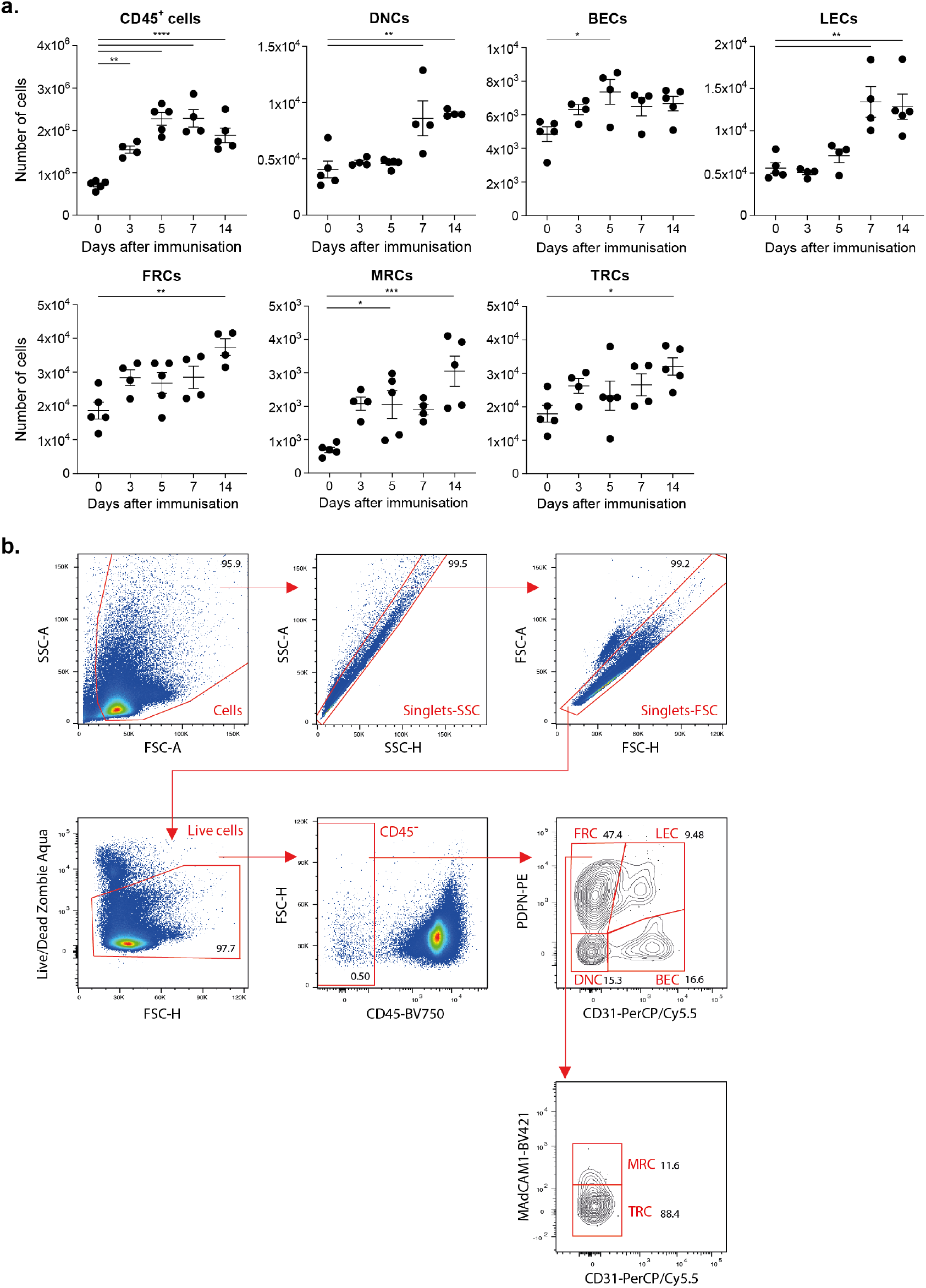
**a.** Analysis by flow cytometry of number of leukocytes (CD45^+^) and indicated stromal cell populations in cell suspensions of inguinal lymph nodes from C57BL/6 mice immunized with IFA/OVA for indicated time points. \**p*<0.05, \*\**p*<0.01, \*\*\**p*<0.001, \*\*\*\**p*<0.0001. **b.** Gating strategy for analysis of *in vivo* immunised lymph nodes. All gates are based on FMOs (fluorescence minus one samples) and relevant controls. Numbers indicate percentage of parental population. BECs = blood endothelial cells, BV = Brilliant Violet, DNCs = double negative cells, FRCs = fibroblastic reticular cells, LECs = lymphatic endothelial cells, MRCs = marginal reticular cells, TRCs = T-zone FRCs.

